# Membrane PI(4,5)P_2_ and ErbB2 abundance regulate ErbB receptor oligomerization and kinase activation

**DOI:** 10.64898/2026.08.24.746611

**Authors:** Mitsuhiro Abe, Masataka Yanagawa, Yasushi Sako

## Abstract

Because ErbB receptors play distinct roles in regulating diverse cellular functions, the mechanisms governing ErbB receptor activation are likely to be more diverse than previously recognized. Phosphatidylinositol 4,5-bisphosphate [PI(4,5)P_2_] positively regulates ErbB1 kinase activity, but the role of PI(4,5)P_2_ in regulating other ErbB family members in living cells remains poorly understood. We show that disruption of PI(4,5)P_2_ binding enhances ErbB4 oligomerization and kinase activity while reducing both processes in ErbB1. Analysis of chimeric receptors identified the juxtamembrane (JM) regions of ErbB1 and ErbB4 as key determinants of their distinct responses to PI(4,5)P_2_ during receptor oligomerization and kinase activation. Furthermore, the JM–kinase module of ErbB1 is more active in the presence of PI(4,5)P_2_, whereas that of ErbB4 is activated by the disruption of PI(4,5)P_2_ binding. In contrast, the JM–kinase module of ErbB2 exhibits weak dependence on PI(4,5)P_2_. ErbB2 preferentially promotes ErbB4 oligomerization over ErbB1 oligomerization, thereby enhancing ErbB4 activation. Collectively, these findings identify plasma membrane PI(4,5)P_2_ availability and ErbB2 abundance as two factors that jointly govern ErbB receptor oligomerization and activation.

## Introduction

The ErbB family of receptor tyrosine kinases (RTKs) plays a central role in regulating cell proliferation, differentiation, and survival through downstream signaling pathways such as Ras/MAPK and PI3K/Akt (Avraham and Yarden, 2011; Lemmon and Schlessinger, 2010; Lemmon et al., 2014). Aberrant regulation of ErbB signaling is a hallmark of many cancers, driving uncontrolled cell proliferation and tumor progression (Yarden and Sliwkowski, 2001). The ErbB family comprises four closely related receptors: ErbB1 (EGFR), ErbB2 (HER2), ErbB3 (HER3), and ErbB4 (HER4) (Kovacs et al., 2015; Lemmon and Schlessinger, 2010; Lemmon et al., 2014). Genetic alterations affecting these receptors, including gene amplification, overexpression, and activating mutations, have been extensively characterized in a wide range of malignancies, particularly lung cancer, breast cancer, and glioma (Bièche et al., 2003; Greulich et al., 2012; Yarden, 2001).

ErbB4 is distinguished by its highly context-dependent biological functions. Although it activates many of the same downstream signaling pathways as the other ErbB receptors, ErbB4 frequently acts as a negative regulator of cell proliferation and its expression is often downregulated or lost in human cancers, consistent with a tumor-suppressive role (Arteaga and Engelman, 2014; Muraoka-Cook et al., 2008; Naresh et al., 2006; Segers et al., 2020). However, in certain tumor contexts, activating mutations or other molecular alterations in ErbB4 promote oncogenic signaling and contribute to tumor development (Prickett et al., 2009; Segers et al., 2020). These seemingly opposing observations suggest that the mechanisms governing ErbB receptor activation and downstream signaling are considerably more diverse than previously appreciated.

Differences in protein function ultimately arise from differences in protein structure. The ErbB family shares a conserved domain architecture consisting of an extracellular region with four subdomains, a single-pass transmembrane helix, a short juxtamembrane (JM) region, a tyrosine kinase domain, and a C-terminal tail (Lemmon and Schlessinger, 2010; Lemmon et al., 2014). Dimerization is essential for the activation of all ErbB receptors. The activation mechanism of ErbB1 is the most extensively characterized of all family members. Binding of epidermal growth factor (EGF) to ErbB1 induces a conformational rearrangement that exposes the extracellular dimerization arm, thereby promoting receptor dimerization. This extracellular conformational change is coupled to association of the N-terminal transmembrane helices and dimerization of the JM region, which together facilitate the formation of the asymmetric intracellular kinase dimer required for catalytic activation (Arkhipov et al., 2013; Ogiso et al., 2002; Red Brewer et al., 2009; Thiel and Carpenter, 2007; Zhang et al., 2006). The activated receptor dimers then undergo autophosphorylation of multiple tyrosine residues within their C-terminal tails, creating docking sites for the adaptor and effector proteins that initiate diverse downstream signaling pathways.

The JM region of ErbB1 is a critical allosteric element that couples ligand binding to kinase activation (Endres et al., 2013; Jura et al., 2009b). It comprises the N-terminal JM-A segment, which forms an antiparallel helical dimer, and the C-terminal JM-B segment, which interacts with the kinase domain (Jura et al., 2009a). These interactions promote the formation and stabilization of the asymmetric kinase dimer, thereby facilitating receptor activation. Phosphatidylinositol 4,5-bisphosphate [PI(4,5)P_2_], a phospholipid enriched in the inner leaflet of the plasma membrane, interacts with basic residues in the JM-A segment and stabilizes its antiparallel dimeric conformation. This stabilization promotes assembly of the active asymmetric kinase dimer and enhances receptor activation (Abd Halim et al., 2015; Hedger et al., 2015; Matsushita et al., 2013; McLaughlin et al., 2005). Consistent with these predictions, our biochemical and single-molecule imaging studies demonstrated that PI(4,5)P_2_ stabilizes the JM-A dimer and promotes ErbB1 oligomerization and activation (Abe et al., 2024; Maeda et al., 2018; Maeda et al., 2022). Although PI(4,5)P_2_-dependent regulation has been established for ErbB1, it is unknown whether this mechanism is conserved in other ErbB family members.

To better understand the diversity of the regulatory mechanisms across the ErbB family members, we investigated the role of PI(4,5)P_2_ in regulating ErbB receptor activation. Because ErbB2 lacks a known ligand (Cho et al., 2003) and ErbB3 exhibits severely impaired kinase activity (Jura et al., 2009b), we focused on ErbB4 to determine whether PI(4,5)P_2_-mediated regulation extends beyond ErbB1. Using mutational analysis, chimeric receptors, single-molecule imaging, and biochemical assays, we demonstrate that PI(4,5)P_2_ has opposite effects on ErbB1 and ErbB4 activation that are mediated by their receptor-specific JM regions. Our findings indicate that PI(4,5)P_2_ stabilizes the active conformation of ErbB1, while restraining ErbB4 activation. Furthermore, ErbB2 expression promotes ErbB4 oligomerization and enhances ErbB4 activation. In summary, this study identified plasma membrane PI(4,5)P_2_ levels and ErbB2 abundance as factors that shape receptor oligomerization and ErbB signaling.

## Results

### PI(4,5)P_2_ binding suppresses ErbB4 kinase activity

We previously showed that substitution of three arginine residues within the ErbB1 JM-A segment with asparagine [ErbB1(3N)] abolishes PI(4,5)P_2_ binding and reduces receptor dimerization, oligomerization, and kinase activity (Abe et al., 2024). The corresponding basic residues are conserved at homologous positions within the ErbB4 JM-A segment (Figure 1A), and molecular dynamics simulations predicted they mediate PI(4,5)P_2_ binding (Hedger et al., 2015). We therefore substituted these three residues with asparagine to generate the PI(4,5)P_2_-binding-deficient mutant ErbB4 [ErbB4(3N)].

**Figure 1.**
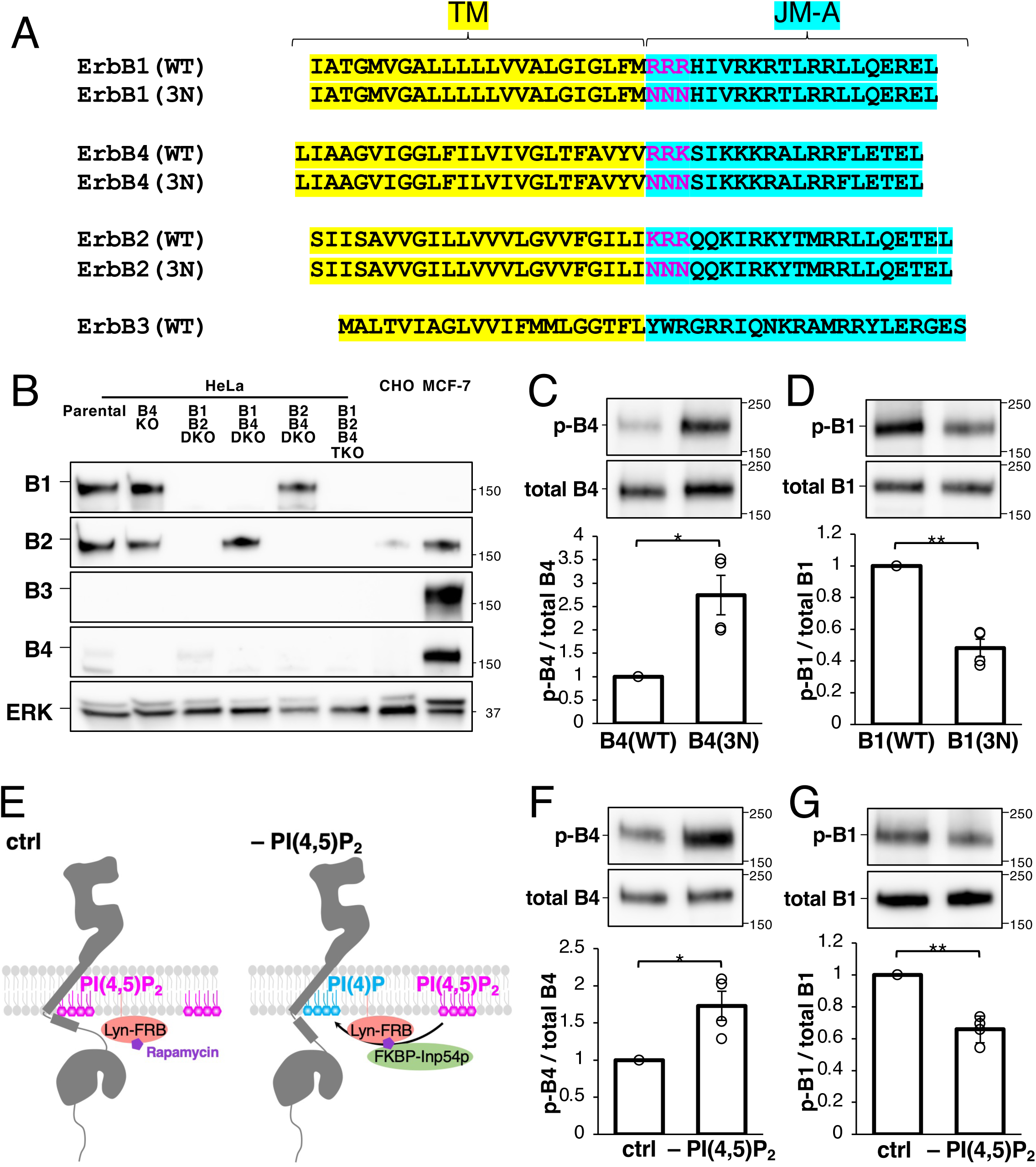
Opposite effects of PI(4,5)P_2_ on ErbB1 and ErbB4 kinase activities. **(A)** Sequence alignment of the transmembrane (TM) and juxtamembrane (JM) regions of ErbB1, ErbB2, ErbB3, and ErbB4. The TM region and JM-A segment are highlighted in yellow and cyan, respectively. Magenta letters indicate the three conserved basic residues in the wild-type (WT) receptors and the corresponding asparagine substitutions in the PI(4,5)P_2_-binding-deficient (3N) mutants. **(B)** Western blot analysis of ErbB family proteins in parental and knockout HeLa cells, CHO-K1 cells, and MCF-7 cells. Cells were lysed, and equal amounts of protein were subjected to Western blot analysis. **(C)** Western blot analysis of ErbB4(WT) and the PI(4,5)P_2_-binding-deficient mutant ErbB4(3N). The ratio of phospho-ErbB4 to total ErbB4 was normalized to that of lane 1. Following overnight starvation, cells were stimulated with 20 nM HRG for 5 min. Data are presented as the mean ± SEM from four independent experiments. **(D)** Western blot analysis of ErbB1(WT) and the PI(4,5)P_2_-binding-deficient mutant ErbB1(3N). The ratio of phospho-ErbB1 to total ErbB1 was normalized to that of lane 1. Following overnight starvation, cells were stimulated with 20 nM EGF for 2 min. Data are presented as the mean ± SEM from four independent experiments. **(E)** Schematic illustration of acute plasma membrane PI(4,5)P_2_ depletion. Cells were transfected with Lyn_11_–FRB alone (left) or together with FKBP–Inp54p (right). Addition of rapamycin recruits FKBP–Inp54p to the plasma membrane, resulting in PI(4,5)P_2_ depletion. **(F)** Western blot analysis of ErbB4 under PI(4,5)P_2_-depleted conditions. The ratio of phospho-ErbB4 to total ErbB4 was normalized to that of lane 1. Data are presented as the mean ± SEM from four independent experiments. **(G)** Western blot analysis of ErbB1 under PI(4,5)P_2_-depleted conditions. The ratio of phospho-ErbB1 to total ErbB1 was normalized to that of lane 1. Data are presented as the mean ± SEM from four independent experiments. *p* < 0.05 (*), *p* < 0.01 (**), *p* < 0.001 (***), and *p* ≥ 0.05 (NS) (t-test).

Heregulin (HRG) stimulation induces both ErbB4 homodimerization and ErbB2–ErbB4 heterodimerization (Ferguson et al., 2000; Okada et al., 2022; Singh et al., 2024; Trenker et al., 2024). To specifically examine the kinase activity of ErbB4 homodimers, we generated ErbB2/ErbB4 double-knockout (DKO) HeLa cells (Figure 1B). As previously reported (Okada et al., 2022), ErbB3 expression was undetectable in HeLa cells, unlike in MCF-7 cells (Figure 1B), indicating that signaling from ErbB3-containing heterodimers is unlikely to contribute to the observed responses in ErbB2/ErbB4 DKO cells. Wild-type ErbB4 [ErbB4(WT)] or ErbB4(3N) was expressed in these cells, and receptor phosphorylation was measured after stimulation with 20 nM HRG for 5 min (Figure 1C). For comparison, the kinase activities of ErbB1(WT) and ErbB1(3N) were analyzed in ErbB1/ErbB2 DKO cells after EGF stimulation (Figure 1D). ErbB1(3N) exhibited reduced kinase activity relative to ErbB1(WT), whereas ErbB4(3N) displayed higher kinase activity than ErbB4(WT), indicating that disruption of PI(4,5)P_2_ binding has opposite effects on ErbB1 and ErbB4.

To confirm these findings independently of mutational analysis, we acutely depleted plasma membrane PI(4,5)P_2_ using a chemically inducible phosphatase recruitment system (Figure 1E) (Suh et al., 2006). Acute PI(4,5)P_2_ depletion increased HRG-stimulated ErbB4 kinase activity (Figure 1F), but decreased EGF-stimulated ErbB1 kinase activity (Figure 1G). Taken together, these results demonstrate that PI(4,5)P_2_ promotes ErbB1 kinase activation but suppresses ErbB4 kinase activation.

### Disruption of PI(4,5)P_2_ binding enhances HRG-induced ErbB4 oligomerization in living cells

Using single-molecule tracking (SMT), we previously demonstrated that disruption of PI(4,5)P_2_ binding destabilizes ErbB1 dimerization and oligomerization, resulting in reduced kinase activity (Abe et al., 2024). Because our kinase assays indicated that PI(4,5)P_2_ binding suppresses ErbB4 activation (Figure 1C,F), we hypothesized that PI(4,5)P_2_ negatively regulates ErbB4 dimerization and oligomerization. To test this possibility, we used SMT to determine the oligomeric state and mobility of ErbB4(WT) and ErbB4(3N).

Halo-tagged ErbB4 (ErbB4–Halo) was expressed at low levels in ErbB2/ErbB4 DKO cells, labeled with SaraFluor 650T (SF650), and imaged by total internal reflection fluorescence microscopy (TIRFM) (Figure 2A, Videos 1 and 2). The single-particle trajectories were classified into three motional states (immobile, slow-mobile, and fast-mobile) using a variational Bayesian hidden Markov model (Hiroshima et al., 2018; Yanagawa and Sako, 2021). Stimulation with 20 nM HRG for 5 min increased the immobile fraction of ErbB4(WT) but decreased the fast-mobile fraction. ErbB4(3N) exhibited greater shifts in these populations (Figure 2B). Consistent with these observations, the mean square displacement (MSD) of ErbB4(3N) at Δ*t* = 0.3 s decreased by 47% in HRG-stimulated cells, whereas that of ErbB4(WT) decreased by only 18% (Figure 2C,D). The MSD analysis also revealed that the diffusion coefficient and confinement length of ErbB4(3N) decreased following HRG stimulation, whereas ErbB4(WT) exhibited more modest changes in both parameters (Figure S1A,B).

**Figure 2.**
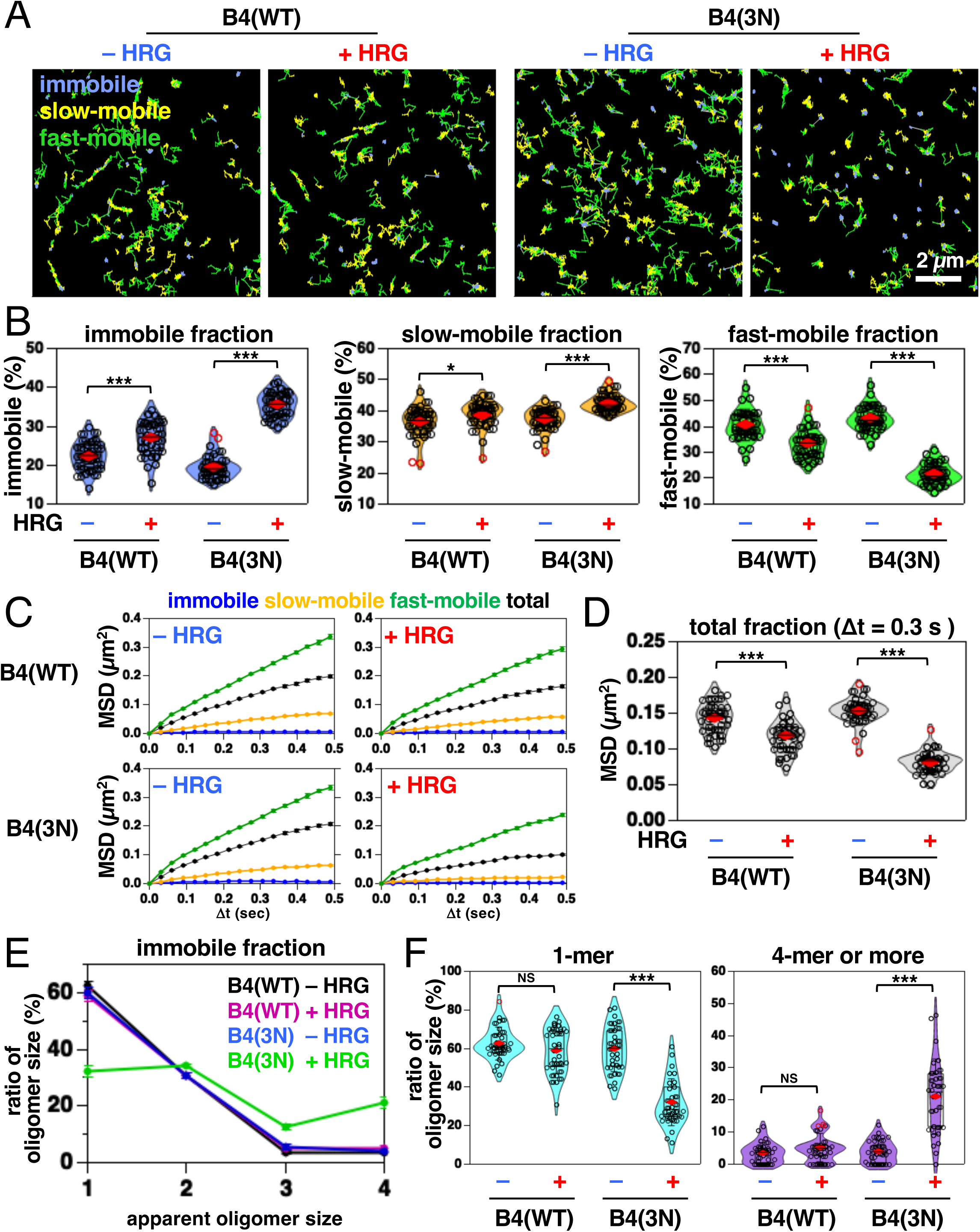
SMT analysis reveals that PI(4,5)P_2_ binding suppresses HRG-induced ErbB4 oligomerization. **(A)** Representative trajectories of ErbB4(WT)–Halo (left) and the PI(4,5)P_2_-binding-deficient mutant ErbB4(3N)–Halo (right) recorded over a 4.5-s observation period. ErbB4(WT)–Halo or ErbB4(3N)–Halo was transiently expressed in ErbB2/ErbB4 DKO HeLa cells. Following overnight starvation, ErbB4–Halo was labeled with the SF650 HaloTag ligand. SMT was performed at a temporal resolution of 30 ms for 4.5 s before HRG stimulation. The same cells were then reimaged 5 min after stimulation with 20 nM HRG. Single-particle trajectories were classified into immobile (blue), slow-mobile (yellow), and fast-mobile (green) fractions. **(B)** Proportions of the immobile (blue), slow-mobile (yellow), and fast-mobile (green) fractions before and after HRG stimulation. Violin plots show the distribution of values from 30 cells. Each circle represents the mean value for an individual cell. **(C)** MSD–Δt plots of ErbB4(WT)–Halo (upper) and ErbB4(3N)–Halo (lower) trajectories before (left) and after (right) HRG stimulation. **(D)** MSD of ErbB4(WT)–Halo and ErbB4(3N)–Halo in the total fraction at Δt = 0.3 s before and after HRG stimulation. Violin plots show the distribution of values from 30 cells. Each circle represents the mean value for an individual cell. **(E)** Distribution of apparent oligomer sizes within the immobile fraction. Putative oligomer sizes were estimated by fitting the distribution of total fluorescence intensities with multiple Gaussian functions. Data are presented as mean ± SEM (*n* = 30 cells). **(F)** Fractions of apparent monomers (left) and tetramers or larger oligomers (right) within the immobile fraction. Violin plots show the distribution of values from 30 cells. Each circle represents the mean value for an individual cell. *p* < 0.05 (*), *p* < 0.01 (**), *p* < 0.001 (***), and *p* ≥ 0.05 (NS) (t-test).

We next quantified the apparent oligomeric states of ErbB4–Halo particles. Consistent with our previous observations for EGF-stimulated ErbB1(WT) (Abe et al., 2024), HRG stimulation predominantly increased the apparent oligomer size of ErbB4(3N) within the immobile fraction (Figure 2E), with minor changes in the slow- and fast-mobile fractions (Figure S1C). Before HRG stimulation, 60% ± 2% of ErbB4(3N) particles were monomers, whereas 4% ± 1% were oligomers containing four or more receptor molecules (Figure 2F; mean ± SEM, *n* = 30 cells). Following HRG stimulation, the proportion of monomers decreased to 32% ± 2%, whereas the proportion of oligomers containing four or more receptor molecules increased to 21% ± 2% (mean ± SEM, *n* = 30 cells). In contrast, a small decrease in the proportion of monomers within the immobile fraction was observed in HRG-stimulated ErbB4(WT) (63% ± 2%) compared with unstimulated cells (59% ± 2%, mean ± SEM, *n* = 30 cells). In unstimulated cells, the mean surface molecule densities of ErbB4(WT) and ErbB4(3N) were comparable (Figure S1D), indicating that differences in receptor expression levels are unlikely to account for the observed differences in oligomerization.

Taken together, these results indicate that PI(4,5)P_2_ binding suppresses HRG-induced ErbB4 oligomerization in living cells.

### Chemically induced dimerization promotes oligomerization of ErbB1(WT) and ErbB4(3N) in living cells

To determine whether disruption of PI(4,5)P_2_ binding within the ErbB4 JM region alters HRG binding or extracellular domain-mediated dimerization, and thereby indirectly affects receptor activation, we replaced the extracellular domains of ErbB receptors with FK506-binding protein (FKBP) and induced receptor dimerization using the chemical dimerizer AP20187. This approach allowed us to examine receptor oligomerization and kinase activation independently of ligand-dependent extracellular interactions.

We first validated this system using ErbB1 by replacing its extracellular domain with FKBP and inducing receptor dimerization with AP20187 in ErbB1/ErbB2 DKO cells (Figure 3A). The SMT analysis showed that AP20187 did not induce the oligomerization of FKBP–ErbB1(WT), which lacks a linker between FKBP and the transmembrane helix. In contrast, the insertion of a 15-amino-acid linker [FKBP–GS15–ErbB1(WT)] enabled AP20187-induced oligomerization to a level comparable to that observed in EGF-stimulated ErbB1(WT). In FKBP–GS15–ErbB1(3N), AP20187 stimulation did not significantly increase the proportion of oligomers within the immobile fraction (Figure S2A). Consistent with these observations, AP20187 increased the phosphorylation of FKBP–GS15–ErbB1(WT), but not that of FKBP–GS15–ErbB1(3N), to levels comparable to those of EGF-stimulated ErbB1(WT) (Figure 3B; Figure S2B). These results demonstrate that the chemically induced dimerization of FKBP–GS15–ErbB1(WT) is sufficient to promote receptor oligomerization and kinase activation.

**Figure 3.**
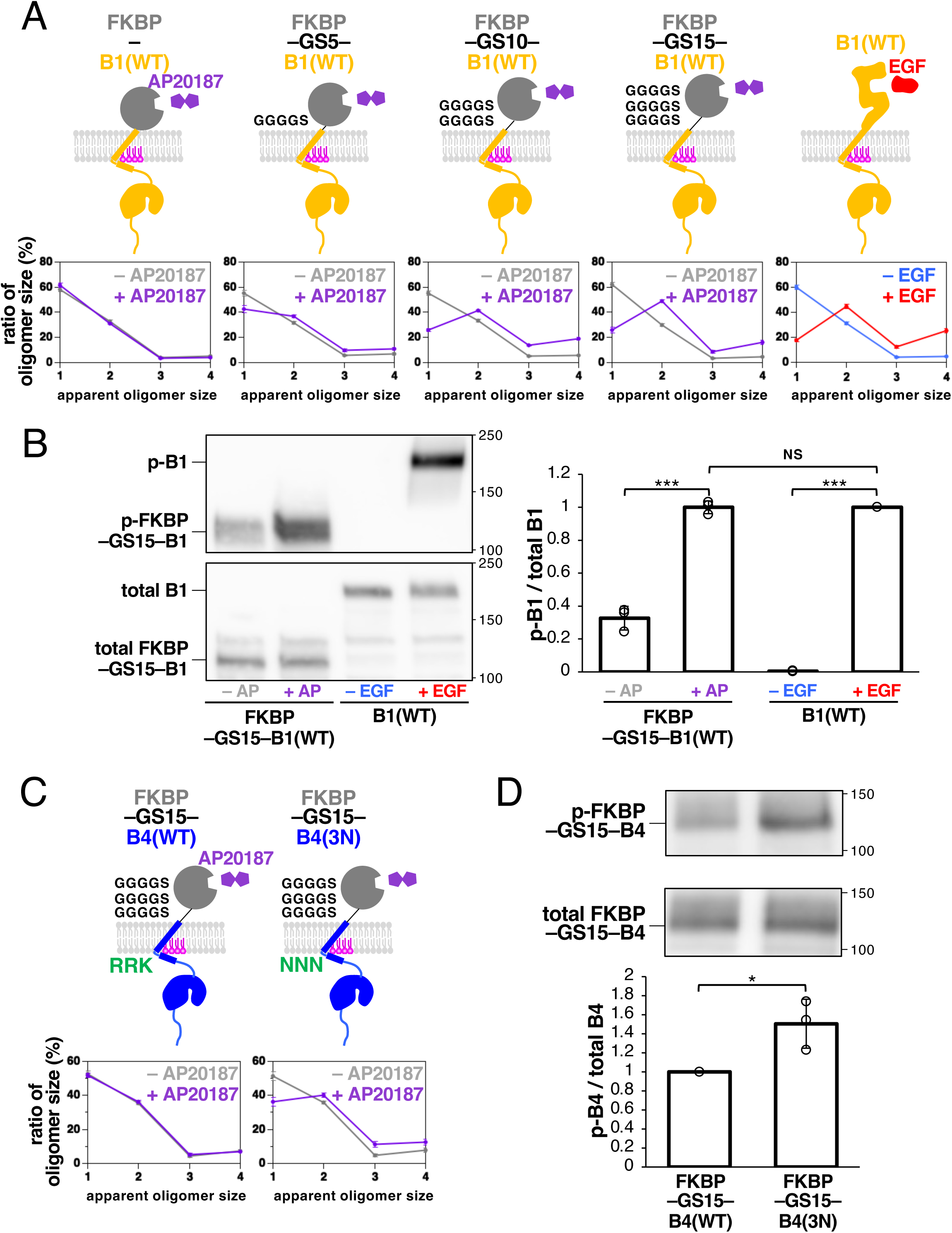
Chemically induced dimerization promotes oligomerization and activation of PI(4,5)P_2_-binding-deficient ErbB4. **(A)** Chemically induced dimerization promotes oligomerization of ErbB1 (upper). The extracellular domain of ErbB1 was replaced with FKBP, and receptor dimerization was induced using the chemical dimerizer AP20187. Four FKBP–ErbB1 constructs were generated with no linker or linkers of 5, 10, or 15 amino acids between FKBP and the transmembrane domain. The distribution of apparent oligomer sizes within the immobile fraction was determined by SMT (lower). Data are presented as the mean ± SEM (*n* = 25–30 cells). **(B)** Western blot analysis of ErbB1 following chemically induced dimerization. The ratio of phospho-ErbB1 to total ErbB1 was normalized to that of lane 4. Data are presented as the mean ± SD from three independent experiments. **(C)** Chemically induced dimerization promotes oligomerization of ErbB4. The extracellular domain of ErbB4 was replaced with FKBP, and receptor dimerization was induced using AP20187. FKBP–GS15–ErbB4(WT) and FKBP–GS15–ErbB4(3N) were analyzed, and the distribution of apparent oligomer sizes within the immobile fraction was determined by SMT. Data are presented as the mean ± SEM (*n* = 30 cells). **(D)** Western blot analysis of ErbB4 following chemically induced dimerization. The ratio of phospho-ErbB4 to total ErbB4 was normalized to that of lane 1. Data are presented as the mean ± SD from three independent experiments. *p* < 0.05 (*), *p* < 0.01 (**), *p* < 0.001 (***), and *p* ≥ 0.05 (NS) (t-test).

We next applied this system to ErbB4 by replacing the ErbB1-derived intracellular region of FKBP–GS15–ErbB1 with the corresponding region of ErbB4 (Figure 3C). The SMT analysis revealed that AP20187 induced the oligomerization of FKBP–GS15–ErbB4(3N), but not that of FKBP–GS15–ErbB4(WT). Similar to HRG-stimulated ErbB4(3N), AP20187 decreased the MSD, increased the immobile fraction, and promoted oligomer formation within the immobile fraction of FKBP–GS15–ErbB4(3N) (Figure S2C-E). In contrast, AP20187 had little effect on FKBP–GS15–ErbB4(WT). Western blotting further showed that AP20187 markedly increased the autophosphorylation of FKBP–GS15–ErbB4(3N), but not that of FKBP–GS15–ErbB4(WT) (Figure 3D).

Taken together, these results demonstrate that the disruption of PI(4,5)P_2_ binding to the ErbB4 JM region enhances receptor oligomerization and kinase activation through mechanisms encoded within the intracellular region following dimerization.

### JM regions encode receptor-specific responses to PI(4,5)P_2_ during ErbB1 and ErbB4 activation

The above results indicate that the disruption of PI(4,5)P_2_ binding within the JM region of ErbB1 reduces receptor dimerization, oligomerization, and kinase activity, whereas the disruption of PI(4,5)P_2_ binding within the corresponding region of ErbB4 enhances these processes. To determine whether the JM regions are responsible for these distinct responses to PI(4,5)P_2_, we generated chimeric receptors in which the JM region of ErbB1 was replaced with that of ErbB4 (Figure 4A). For simplicity, we refer to the original and chimeric receptors according to the origin of their JM regions (i.e., B1JM or B4JM). Receptor oligomerization and kinase activity were then analyzed in ErbB1/ErbB2/ErbB4 triple-knockout (TKO) cells.

**Figure 4.**
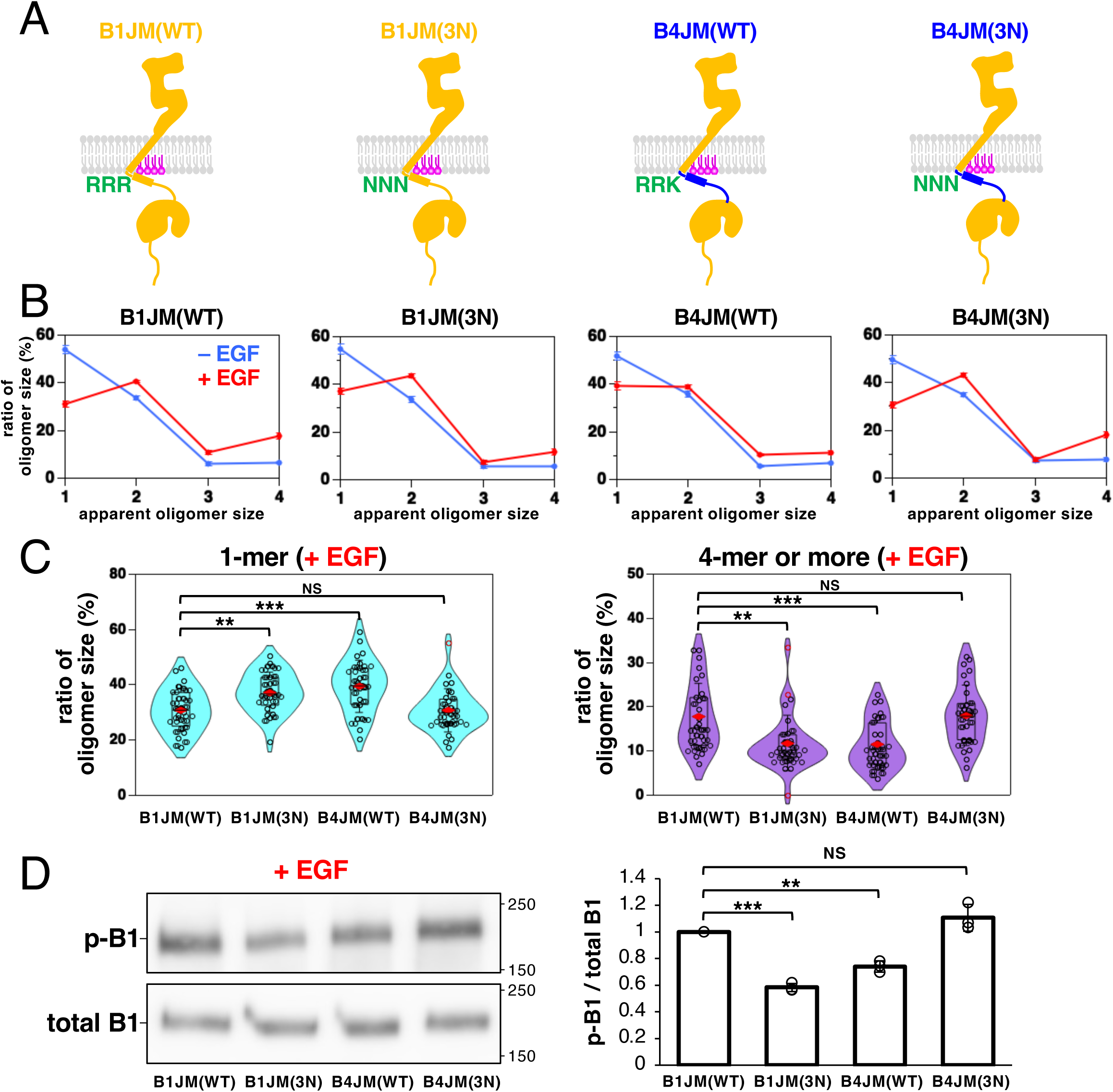
SMT analysis identifies the JM regions of ErbB1 and ErbB4 as determinants of their responses to PI(4,5)P_2_ binding. **(A)** Schematic of the chimeric receptors in which the JM region of ErbB1 was replaced with that of ErbB4. The indicated receptors were transiently expressed in ErbB1/ErbB2/ErbB4 TKO HeLa cells and analyzed by SMT or Western blotting. Following overnight starvation, cells were stimulated with 20 nM EGF for 2 min. **(B)** Distribution of apparent oligomer sizes within the immobile fraction before (blue) and after (red) EGF stimulation (lower). Data are presented as the mean ± SEM (*n* = 30 cells). **(C)** Fractions of apparent monomers (left) and tetramers or larger oligomers (right) within the immobile fraction after EGF stimulation. Violin plots show the distribution of values from 30 cells. Each circle represents the mean value for an individual cell. **(D)** Western blot analysis of the indicated receptors. The ratio of phospho-receptor to total receptor was normalized to that of lane 1. Data are presented as the mean ± SD from three independent experiments. *p* < 0.05 (*), *p* < 0.01 (**), *p* < 0.001 (***), and *p* ≥ 0.05 (NS) (t-test).

The SMT analysis revealed that the receptors could be classified into two groups based on their responses to EGF stimulation (Figure 4B,C). Pronounced increases in oligomerization were observed following stimulation in the first group, consisting of B1JM(WT) and B4JM(3N). In contrast, only modest changes in these parameters were observed in the second group, consisting of B1JM(3N) and B4JM(WT). In the first group, monomers and oligomers containing four or more receptor molecules accounted for approximately 31% and 18% of the immobile fraction, respectively, after EGF stimulation (Figure 4C; *n* = 30 cells). In the second group, these proportions were approximately 39% and 11%, respectively (*n* = 30 cells).

Consistent with the SMT analysis, EGF stimulation induced robust autophosphorylation of B1JM(WT) and B4JM(3N), whereas phosphorylation of B1JM(3N) and B4JM(WT) remained substantially lower (Figure 4D). Taken together, these results indicate that the JM regions of ErbB1 and ErbB4 determine the receptor-specific effects of PI(4,5)P_2_ on receptor oligomerization and kinase activation.

### ErbB1 responds to PI(4,5)P_2_ binding, whereas ErbB4 is activated by the disruption of PI(4,5)P_2_ binding, and ErbB2 exhibits weak dependence on PI(4,5)P_2_

The ErbB family members share a common signaling mechanism but exhibit distinct signaling properties. Because differences in relative kinase activity among ErbB receptors may contribute to their distinct downstream signaling outputs, we compared the kinase activities of receptors containing wild-type or PI(4,5)P_2_-binding-deficient JM regions derived from ErbB1, ErbB2, and ErbB4. To enable a direct comparison under identical stimulation conditions, the extracellular regions of ErbB2 and ErbB4 were replaced with that of ErbB1, allowing activation of all receptors by EGF. In addition, because differences in the C-terminal tail length and phosphorylation sites could influence receptor phosphorylation and SMT analyses, the C-terminal tails of ErbB2 and ErbB4 were replaced with that of ErbB1. To monitor kinase activity using a common phosphorylation site, a 12-amino-acid sequence containing an ErbB3-derived tyrosine phosphorylation site (B3Y) was inserted between the receptor C-terminal tail and a Halo tag, yielding chimeric receptors (Figure 5A). For simplicity, the original and chimeric receptors are referred to according to the origins of their JM region and kinase domain (e.g., B1JMKD or B2JMKD). These constructs were expressed in ErbB1/ErbB2/ErbB4 TKO cells.

**Figure 5.**
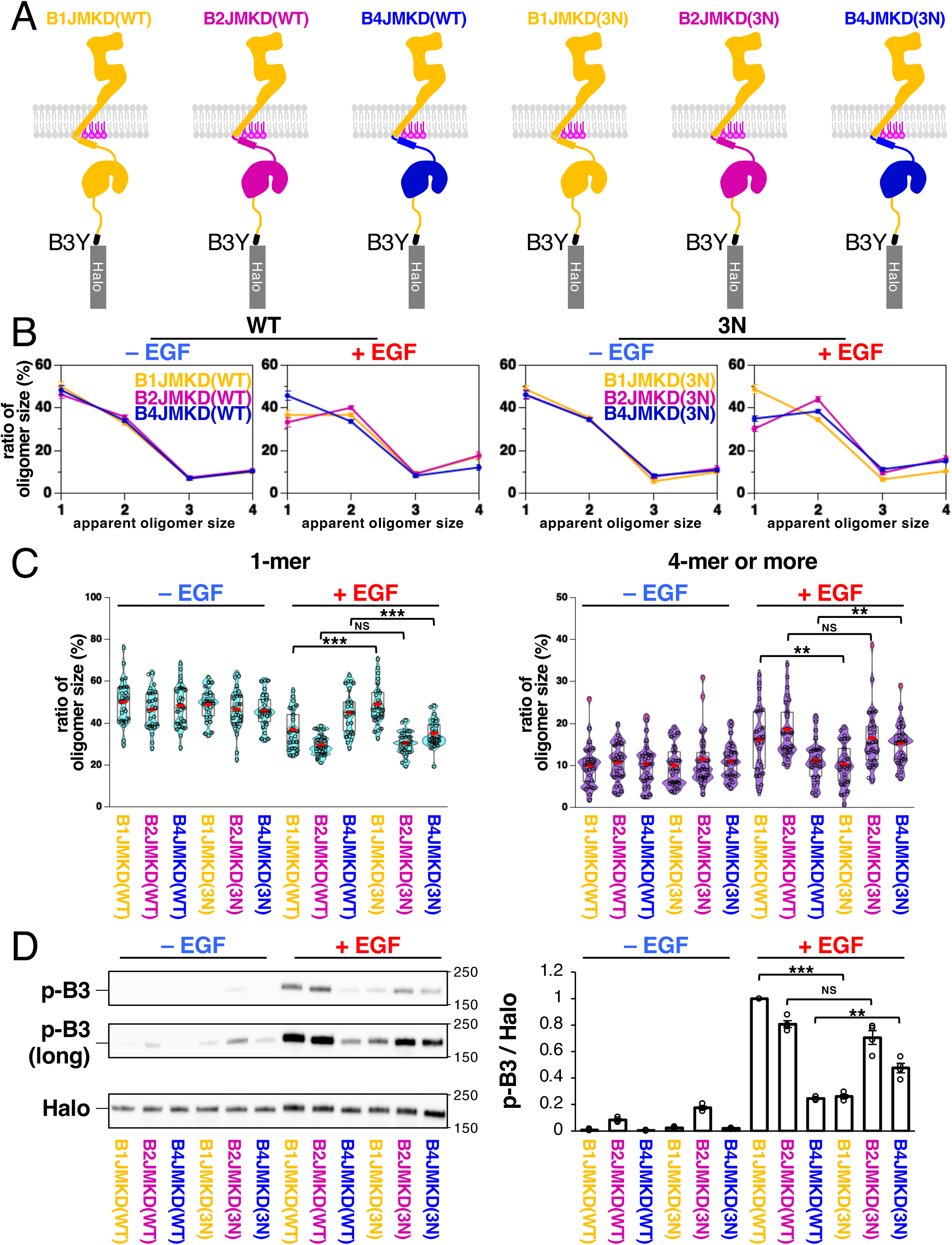
Comparative analysis of PI(4,5)P_2_ responses of ErbB1, ErbB2, and ErbB4. **(A)** Schematic of the B1JMKD, B2JMKD, and B4JMKD chimeric receptor constructs. The JM region and kinase domain (collectively referred to as JMKD) of ErbB1 were replaced with the corresponding regions of ErbB2 or ErbB4. A 12-amino-acid sequence containing an ErbB3-derived tyrosine phosphorylation site (B3Y) was inserted between the receptor C-terminal tail and the Halo tag. **(B)** Distribution of apparent oligomer sizes within the immobile fraction. The indicated chimeric receptors were transiently expressed in ErbB1/ErbB2/ErbB4 TKO HeLa cells. Following transfection, cells were cultured in complete medium for at least 24 h, starved for 3 h, and stimulated with 20 nM EGF for 2 min. Data are presented as the mean ± SEM (*n* = 25 cells). **(C)** Fractions of apparent monomers (left) and tetramers or larger oligomers (right) within the immobile fraction. Violin plots show the distribution of values from 25 cells. Each circle represents the mean value for an individual cell. **(D)** Western blot analysis of the indicated chimeric receptors. The receptors were transiently expressed in ErbB1/ErbB2/ErbB4 TKO HeLa cells. Following transfection, cells were cultured in complete medium for at least 24 h, starved for 3 h, and stimulated with 20 nM EGF for 2 min. Phosphorylated and total receptors were detected using anti-phospho-ErbB3 and anti-Halo antibodies, respectively (left). The ratio of phosphorylated to total receptor was normalized to that of lane 7 (right). Data are presented as the mean ± SEM from four independent experiments. *p* < 0.05 (*), *p* < 0.01 (**), *p* < 0.001 (***), and *p* ≥ 0.05 (NS) (t-test).

We first analyzed receptor oligomerization using SMT (Figure 5B,C; Figure S3A). Because ErbB2 also contains three conserved basic residues within the JM-A segment that are predicted to mediate PI(4,5)P_2_ binding (Hedger et al., 2015), these residues were substituted with asparagine to generate a PI(4,5)P_2_-binding-deficient mutant. Among the receptors examined, EGF stimulation markedly increased the oligomer formation of B1JMKD(WT) within the immobile fraction, whereas B1JMKD(3N) exhibited a weaker oligomerization response (Figure 5C, left). In contrast, EGF stimulation preferentially decreased the oligomer formation of B4JMKD(WT) relative to B4JMKD(3N). By comparison, B2JMKD(WT) and B2JMKD(3N) exhibited similar levels of oligomer formation within the immobile fraction following EGF stimulation. Consistent with these observations, the reduction in MSD at Δ*t* = 0.3 s in EGF-stimulated cells was correlated with the degree of receptor oligomerization (Figure S3B).

To confirm these observations biochemically, we measured receptor autophosphorylation using an anti-phospho-ErbB3 antibody, which recognizes the inserted B3Y sequence (Figure 5D). The antibody did not recognize the receptor lacking the B3Y sequence (Figure S3C). Before EGF stimulation, basal autophosphorylation was greater in the PI(4,5)P_2_-binding-deficient B1JMKD and B4JMKD mutants than in their PI(4,5)P_2_-binding-competent counterparts (Figure S3D). Furthermore, B2JMKD exhibited greater basal activity than B1JMKD and B4JMKD. After EGF stimulation, B1JMKD(WT) exhibited greater autophosphorylation than B1JMKD(3N), whereas B4JMKD(WT) exhibited less autophosphorylation than B4JMKD(3N). In contrast, B2JMKD(WT) and B2JMKD(3N) exhibited comparable levels of autophosphorylation following EGF stimulation. These biochemical results are consistent with the SMT analysis of receptor oligomerization (Figure 5B,C).

Taken together, these results indicate that the JM region, specifically the kinase domain, of ErbB1 is preferentially activated by PI(4,5)P_2_ binding, whereas that of ErbB4 is preferentially activated by the loss of PI(4,5)P_2_ binding. In contrast, the activation of ErbB2 is relatively insensitive to PI(4,5)P_2_.

### ErbB2 preferentially promotes ErbB4 oligomerization over ErbB1 oligomerization and enhances ErbB4 activation

Because the above experiments only examined homodimeric receptors, we next investigated heterodimer formation. In cells expressing ErbB2, HRG stimulation of ErbB4 induces ErbB4–ErbB4 homodimerization or ErbB2–ErbB4 heterodimerization (Ferguson et al., 2000; Trenker et al., 2024). To determine whether ErbB2–ErbB4 heterodimers form in living cells, we analyzed the ErbB2–ErbB4 association by SMT. The cytoplasmic tail of ErbB2 was tagged with SNAP2 (ErbB2–SNAP2) (Kühn et al., 2025), and either ErbB2(WT)–SNAP2 or ErbB2(3N)–SNAP2 was coexpressed with ErbB4(WT)–Halo or ErbB4(3N)–Halo, respectively, in ErbB2/ErbB4 DKO cells at low expression levels. ErbB2–SNAP2 and ErbB4–Halo were labeled with SNAP-Cell TMR-Star and SF650, respectively, and analyzed by TIRFM (Figure 6A, Videos 3 and 4).

**Figure 6.**
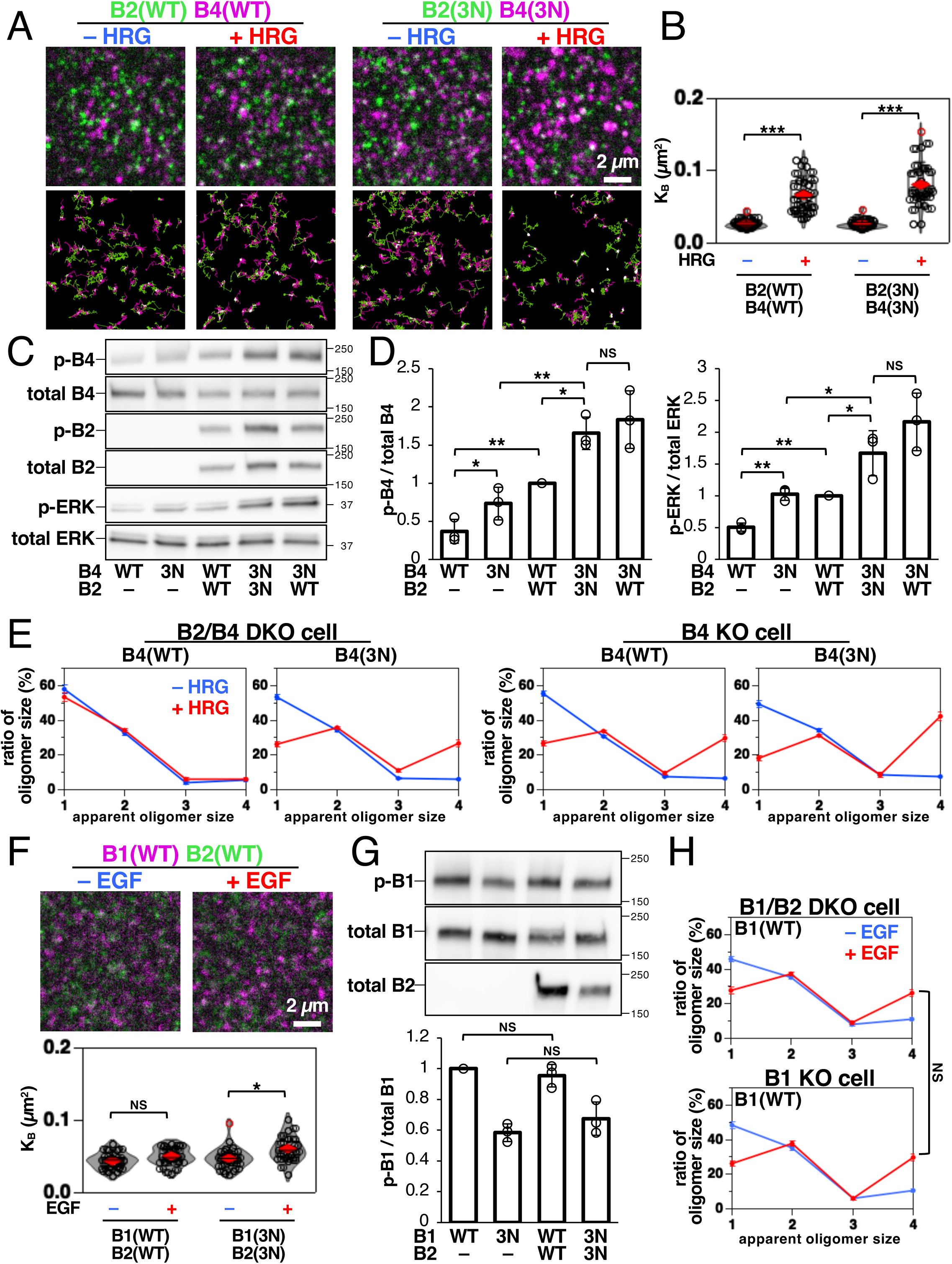
ErbB2 promotes ErbB4 association and enhances ErbB4 kinase activity following HRG stimulation. **(A)** SMT analysis of ErbB2–ErbB4 association. Representative TIRFM images (upper) and corresponding trajectories (lower). ErbB2(WT)–SNAP2 and ErbB4(WT)–Halo, or ErbB2(3N)–SNAP2 and ErbB4(3N)–Halo, were coexpressed in ErbB2/ErbB4 DKO cells at low expression levels. ErbB2–SNAP2 (green) and ErbB4–Halo (magenta) were labeled with the respective ligands. Scale bar, 2 µm. **(B)** Colocalization of ErbB2 and ErbB4 before and after HRG stimulation. K_B_ was defined as the density of colocalized ErbB2–ErbB4 particles normalized to the product of the densities of free ErbB2 and ErbB4 particles. Violin plots show the distribution of values from 30 cells. Each circle represents the mean value for an individual cell. **(C)** Western blot analysis of ErbB4 and ERK phosphorylation in the absence or presence of ErbB2. ErbB4 alone or ErbB2 together with ErbB4 was expressed in ErbB2/ErbB4 DKO cells. Following transfection, cells were cultured in complete medium for 24 h, starved for 6 h, and stimulated with 20 nM HRG for 30 min. **(D)** Quantification of phospho-ErbB4/total ErbB4 (left) and phospho-ERK/total ERK (right). Values were normalized to those of lane 3. Data are presented as the mean ± SD from three independent experiments. **(E)** Distribution of apparent oligomer sizes of ErbB4(WT) and ErbB4(3N) within the immobile fraction. ErbB4(WT) or ErbB4(3N) was expressed in ErbB2/ErbB4 DKO cells (left) or ErbB4 KO cells expressing ErbB2 (right), and receptor oligomerization was analyzed by SMT before (blue) and after (red) HRG stimulation. Data are presented as the mean ± SEM (*n* = 30 cells). **(F)** SMT analysis of ErbB1–ErbB2 association. Representative TIRFM images (upper) and K_B_ values (lower). ErbB1(WT)–Halo and ErbB2(WT)–SNAP2, or ErbB1(3N)–Halo and ErbB2(3N)–SNAP2, were coexpressed in ErbB1/ErbB2 DKO cells at low expression levels. ErbB1–Halo (magenta) and ErbB2–SNAP2 (green) were labeled with the respective ligands. Violin plots show the distribution of values from 25 cells. Each circle represents the mean value for an individual cell. **(G)** Western blot analysis of ErbB1 phosphorylation in the absence or presence of ErbB2. ErbB1 alone or ErbB2 together with ErbB1 was expressed in ErbB1/ErbB2 DKO cells. Following transfection, cells were cultured in complete medium for 24 h, starved for 6 h, and stimulated with 20 nM EGF for 30 min. The ratio of phospho-ErbB1 to total ErbB1 was normalized to that of lane 1. Data are presented as the mean ± SD from three independent experiments. **(H)** Distribution of apparent oligomer sizes of ErbB1(WT) within the immobile fraction. ErbB1(WT) was expressed in ErbB1/ErbB2 DKO cells (upper) or ErbB1 KO cells expressing ErbB2 (lower), and receptor oligomerization was analyzed by SMT. Data are presented as the mean ± SEM (*n* = 25 cells). *p* < 0.05 (*), *p* < 0.01 (**), *p* < 0.001 (***), and *p* ≥ 0.05 (NS) (t-test).

Consistent with the results described above (Figure 2E,F), ErbB4(3N) exhibited larger changes in apparent oligomer size after HRG stimulation compared with ErbB4(WT). In contrast, ErbB2 oligomerization did not increase following HRG stimulation (Figure S4A). To quantify the ErbB2–ErbB4 association, we calculated the apparent binding affinity (K_B_) as the density of colocalized ErbB2–ErbB4 particles normalized to the product of the densities of free ErbB2 and ErbB4 particles (Kuwashima et al., 2024). HRG stimulation increased K_B_ for both the ErbB2(WT)/ErbB4(WT) and ErbB2(3N)/ErbB4(3N) pairs, indicating that HRG promotes the ErbB2–ErbB4 association.

To determine whether this association was specific to ErbB2, we next examined ErbB1–ErbB4 interactions using ErbB1 tagged with mStayGold (ErbB1–mSG) (Ando et al., 2024). ErbB1(WT)–mSG or ErbB1(3N)–mSG was coexpressed with ErbB4(WT)–Halo or ErbB4(3N)–Halo in ErbB1/ErbB4 DKO cells. In contrast to the ErbB2–ErbB4 pairs, HRG stimulation did not increase the K_B_ between ErbB1 and ErbB4 (Figure S4B,C). These results indicate that HRG preferentially promotes the association of ErbB4 with ErbB2, but not with ErbB1, under our experimental conditions.

We next examined the effect of ErbB2 on ErbB4 kinase activity (Figure 6C,D). ErbB4(WT) alone or together with ErbB2(WT) was expressed in ErbB2/ErbB4 DKO cells under stimulation with HRG. Measurement of ErbB4 phosphorylation showed that ErbB2 expression enhanced ErbB4 kinase activity. Consistent with this result, ERK phosphorylation was also increased in the presence of ErbB2. We next examined the effect of disrupting PI(4,5)P_2_ binding in both receptors. Similar to the results described above (Figure 1C), ErbB4(3N) exhibited greater kinase activity than ErbB4(WT). Coexpression of ErbB2(3N) with ErbB4(3N) further increased ErbB4 phosphorylation beyond that observed in the ErbB2(WT)/ErbB4(WT) pair. This enhancement was not due to intrinsically greater ErbB2(3N) activity, because the coexpression of ErbB2(WT) also increased ErbB4(3N) phosphorylation. These results indicate that ErbB2 enhances ErbB4 kinase activity and that disruption of PI(4,5)P_2_ binding to ErbB4 further potentiates this effect.

Because ErbB2 enhanced ErbB4 phosphorylation, we next examined whether it also promotes ErbB4 oligomerization after HRG stimulation. ErbB4(WT) or ErbB4(3N) was expressed in ErbB4-knockout (KO) cells highly expressing ErbB2 (Figure 1B), and receptor oligomerization was analyzed by SMT (Figure 6E). Following HRG stimulation, ErbB4 oligomerization showed a greater increase in ErbB2-expressing cells than in ErbB2/ErbB4 DKO cells. In ErbB2-expressing cells, the proportion of monomers decreased to 18 ± 2% ErbB4(3N), but the proportion of oligomers containing four or more receptor molecules increased to 42 ± 3% (mean ± SEM, *n* = 30 cells) (Figure 6, Figure S4D).

ErbB4(WT) also exhibited enhanced oligomerization; the proportion of monomers decreased to 27 ± 2% and that that of oligomers containing four or more receptor molecules increased to 30 ± 2% (mean ± SEM, *n* = 30 cells). Consistent with these findings, both ErbB4(3N) and ErbB4(WT) exhibited greater changes in MSD in HRG-stimulated ErbB2-expressing cells than in ErbB2/ErbB4 DKO cells (Figure S4E). These results show that ErbB2 promotes ErbB4 oligomerization and enhances ErbB4 activation, particularly when PI(4,5)P_2_ binding to ErbB4 is disrupted.

We next examined whether ErbB2 promotes ErbB1 oligomerization and enhances ErbB1 activation. The SMT analysis showed that EGF stimulation produced a smaller increase in ErbB1–ErbB2 association relative to the effect of HGF stimulation on the ErbB2–ErbB4 association under our experimental conditions (Figure 6F). This result is consistent with previous findings that only a small fraction of ErbB1–ErbB2 heterodimers can be detected by live-cell single-molecule imaging (Bai et al., 2023) and that ErbB2–ErbB4 heterodimers appear to be the most stable of ErbB2-containing heterodimers (Trenker et al., 2024). We then assessed the effect of ErbB2 expression on ErbB1 kinase activity (Figure 6G). Measurement of ErbB1 phosphorylation showed that ErbB2 expression did not enhance ErbB1 kinase activity. Consistent with these findings, ErbB2 expression did not markedly increase ErbB1 oligomerization following EGF stimulation, because ErbB1 oligomerization was comparable between ErbB1 KO cells expressing ErbB2 and ErbB1/ErbB2 DKO cells (Figure 6H). These results indicate that ErbB2 preferentially promotes ErbB4 oligomerization over ErbB1 oligomerization, thereby enhancing ErbB4 activation.

## Discussion

Here, we investigated the role of PI(4,5)P_2_ in regulating ErbB receptor activation. Our results suggest that PI(4,5)P_2_ binding promotes ErbB1 activation, whereas the disruption of PI(4,5)P_2_ binding promotes ErbB4 activation. In contrast, ErbB2 exhibits relatively weak dependence on PI(4,5)P_2_ for activation. Although ErbB receptors activate many common downstream pathways, including Ras/MAPK, PI3K/Akt, PLCγ, and Src/FAK signaling, each ErbB receptor exhibits distinct signaling preferences that give rise to diverse biological outputs (Citri and Yarden, 2006; Hynes and Lane, 2005; Lemmon and Schlessinger, 2010; Yarden and Sliwkowski, 2001). ErbB1 preferentially activates PLCγ signaling over the PI3K/Akt pathway in HeLa cells (Abe et al., 2024), whereas ErbB4 favors the PI3K/Akt and STAT5 pathways (Brockhoff, 2022; Jones et al., 1999; Telesco et al., 2013). Based on these findings, we propose a model in which ErbB1–ErbB1 and ErbB1–ErbB2 dimers are preferentially activated under conditions of high PI(4,5)P_2_ availability, whereas ErbB2–ErbB4 and ErbB4–ErbB4 dimers are favored when PI(4,5)P_2_ availability is reduced (Figure 7, upper). Thus, changes in plasma membrane PI(4,5)P_2_ availability may dynamically reshape the ErbB receptor activation landscape by promoting the formation of specific receptor oligomers with distinct signaling outputs.

**Figure 7.**
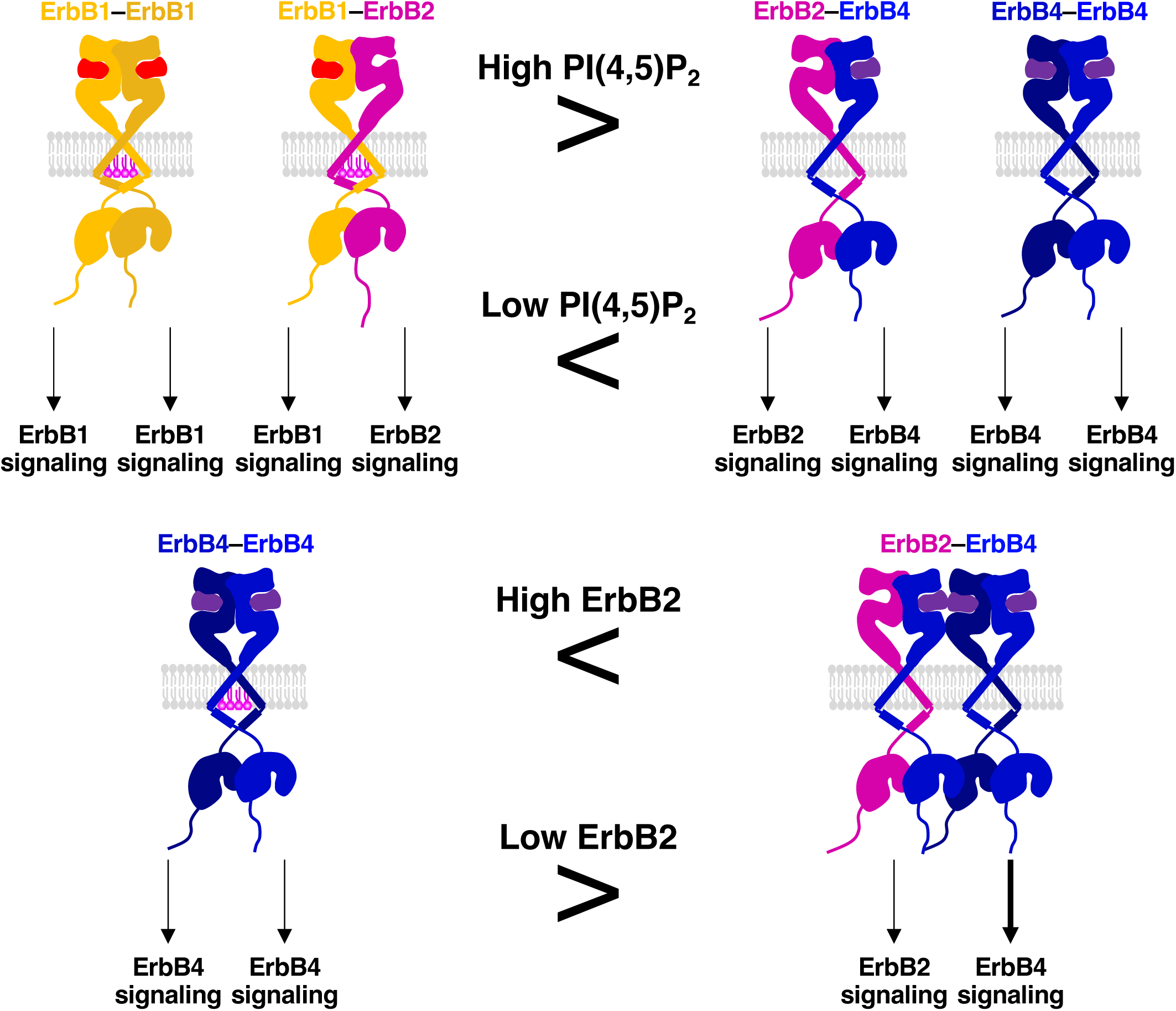
Proposed model illustrating how plasma membrane PI(4,5)P_2_ availability and ErbB2 abundance shape ErbB receptor oligomerization and signaling. **(A)** Model illustrating the contribution of plasma membrane PI(4,5)P_2_ availability to ErbB receptor activation. ErbB1–ErbB1 and ErbB1–ErbB2 complexes are preferentially activated under conditions of high PI(4,5)P_2_ availability, whereas ErbB4-containing complexes are preferentially activated when PI(4,5)P_2_ availability is reduced. **(B)** Model illustrating the contribution of ErbB2 abundance to ErbB4 oligomerization and activation. Increased ErbB2 expression promotes ErbB4 oligomerization and enhances ErbB4 activity, potentially through the formation of ErbB2–ErbB4 heterodimers, thereby influencing downstream signaling.

In addition to PI(4,5)P_2_, the level of ErbB2 expression represents another important determinant of downstream signaling pathway selection. ErbB2 expression varies substantially among cell types and tissues in physiologic and pathologic conditions (Hynes and Lane, 2005; Slamon et al., 1987; Yarden and Sliwkowski, 2001). Because our data suggest that ErbB2 preferentially promotes ErbB4 oligomerization over ErbB1 oligomerization (Figure 6E–H), increased ErbB2 expression would be expected to enhance ErbB4 oligomerization (Figure 7, lower). However, our SMT analysis showed that ErbB2 oligomerization did not increase when subjected to the conditions that enhanced ErbB4 oligomerization following HRG stimulation (Figure S4A). This suggests that ErbB2 may promote ErbB4 oligomerization without being extensively incorporated into the resulting ErbB4 oligomers. The mechanism underlying ErbB2-dependent ErbB4 oligomerization remains unclear. ErbB2 has been implicated in PLCγ1 signaling, because activated ErbB2 was reported to promote PLCγ1 tyrosine phosphorylation (Peles et al., 1991). One possible mechanism is that, following activation of ErbB2–ErbB4 heterodimers, ErbB2 promotes local PI(4,5)P_2_ depletion through PLCγ1, which may in turn modulate ErbB4 oligomerization. Although this model suggests that ErbB2 abundance and PI(4,5)P_2_ availability may be mechanistically coupled, alternative mechanisms cannot be excluded. Thus, membrane PI(4,5)P_2_ levels and ErbB2 abundance might both influence the selection of ErbB signaling pathways and modulate the persistence of receptor-mediated signaling. Future quantitative phosphoproteomic and proteomic analyses under experimental conditions aimed at manipulating PI(4,5)P_2_ levels and ErbB2 expression will help define how these factors influence the selection and dynamics of the ErbB signaling networks.

Upon EGF stimulation, the formation of an antiparallel JM-A dimer between two ErbB1 molecules promotes the formation and stabilization of the asymmetric kinase dimer, thereby facilitating receptor activation (Endres et al., 2013; Jura et al., 2009b). Molecular dynamics simulations indicate that PI(4,5)P_2_ interacts with basic residues within the JM-A segment, stabilizing the antiparallel JM-A dimer of ErbB1 (Abd Halim et al., 2015). Consistent with this model, AlphaFold3 predictions using two ErbB1 JM-A peptides did not generate an antiparallel dimer in the absence of PI(4,5)P_2_, suggesting that PI(4,5)P_2_ binding may be important for stabilizing this conformation (Figure S5). In contrast, AlphaFold3 predicted that two ErbB4 JM-A peptides readily form an antiparallel dimer in the absence of PI(4,5)P_2_, implying that PI(4,5)P_2_ binding is less critical for antiparallel JM-A dimer formation in ErbB4 than in ErbB1. This structural difference may explain why ErbB4 is preferentially activated under PI(4,5)P_2_-binding-deficient or PI(4,5)P_2_-depleted conditions following HRG stimulation. However, this model does not explain why ErbB4 is not fully activated in the presence of PI(4,5)P_2_. Future molecular dynamics simulations of the ErbB4 JM-A segment in the presence and absence of PI(4,5)P_2_ should help clarify how PI(4,5)P_2_ influences the stability and dynamics of the antiparallel JM-A dimer and, consequently, ErbB4 activation.

Interestingly, basal autophosphorylation before EGF stimulation was greater in the PI(4,5)P_2_-binding-deficient B1JMKD and B4JMKD mutants than in their PI(4,5)P_2_-binding-competent counterparts (Figure S3D). These results suggest that PI(4,5)P_2_ suppresses the basal activity of both ErbB1 and ErbB4. Increased basal autophosphorylation of ErbB1 has also been reported under cholesterol-depleted conditions (Kim et al., 2025; Takayama et al., 2024). Likewise, the disruption of tetraspanin CD82 increased basal ErbB1 activity before ligand stimulation (Restrepo Cruz et al., 2026). Cholesterol was shown to bind to CD82 (Huang et al., 2020) and promote the clustering of PI(4,5)P_2_ (Lolicato et al., 2022). These observations raise the possibility that specialized plasma membrane domains enriched in cholesterol and PI(4,5)P_2_ clusters may contribute to the suppression of ligand-independent activation of ErbB1 and ErbB4. Such spatial regulation may help maintain receptors in an inactive state until ligand stimulation occurs. In contrast to the B1JMKD and B4JMKD chimeras, the B2JMKD chimera exhibited elevated basal activity irrespective of its PI(4,5)P_2_-binding status. This observation is consistent with the previous findings that ErbB2 readily forms ligand-independent homodimers when overexpressed (Ma et al., 2026) and that ErbB2 is frequently overexpressed in breast cancer, where spontaneous receptor activation contributes to oncogenic signaling (Atallah et al., 2025; Moasser, 2007; Slamon et al., 1987). Taken together, these findings suggest that the mechanisms that restrict basal receptor activity differ among the ErbB family members and may reflect their distinct physiologic and pathologic functions.

In conclusion, we have identified PI(4,5)P_2_ as a molecular switch that differentially controls the activation and signaling properties of ErbB family members. By modulating receptor oligomerization and kinase activation in a receptor-specific manner, PI(4,5)P_2_ contributes to the diverse ErbB signaling outputs. Given that putative PI(4,5)P_2_-binding residues are predicted to be conserved in many RTKs (Hedger et al., 2015), similar lipid-dependent regulatory mechanisms may operate more broadly across the RTK superfamily. Future studies combining quantitative lipidomics, phosphoproteomics, and biochemical analyses of the local membrane lipid environment surrounding ErbB receptors are needed to understand how membrane lipids coordinate RTK activation and downstream signaling in living cells.

## Materials and Methods

### Cell culture, transfection, and chemical treatments

HeLa cells (#RCB0007) were obtained from the RIKEN BioResource Research Center (Tsukuba, Japan). Plasmids were transiently transfected using the Neon Transfection System (Thermo Fisher Scientific) according to the manufacturer’s instructions. Cells were cultured at 37°C in Dulbecco’s modified Eagle’s medium (DMEM; Nacalai Tesque Inc.) supplemented with 10% fetal bovine serum (FBS).

To minimize receptor endocytosis, all ligand stimulation and chemically induced dimerization experiments were performed at 25°C. Before stimulation, cells were incubated in DMEM containing 0.2% bovine serum albumin (BSA) for at least 3 h at 37°C and then equilibrated at 25°C for at least 10 min. Ligand stimulation was performed with 20 nM epidermal growth factor (EGF; #315-09, PeproTech) or 20 nM heregulin (HRG; #396-HB, R&D Systems). For chemically induced dimerization experiments, AP20187 (#S8487, Selleck Chemicals) was added to a final concentration of 5 μM, and cells were maintained at 25°C throughout the experiment.

### Plasmid construction

pCMV–ErbB2–Halo and pCMV–ErbB4–Halo were generated by replacing the ErbB1 coding sequence in pCMV–EGFR(WT)–Halo (Abe et al., 2024; #RDB20672, RIKEN BRC) with the corresponding coding sequences of ErbB2 and ErbB4 (JM-a CYT-1 isoform), respectively. pCMV–ErbB2(3N)–Halo was generated by introducing the K654N/R655N/R656N substitutions into pCMV–ErbB2–Halo, and pCMV–ErbB4(3N)–Halo was generated by introducing the R651N/R652N/K653N substitutions into pCMV–ErbB4–Halo.

To generate the B2JMKD and B4JMKD chimeras, the coding sequences encoding the JM region and kinase domain of ErbB1 were replaced with the corresponding sequences from ErbB2 and ErbB4, respectively. A 36-bp sequence encoding a 12-amino-acid peptide containing a tyrosine residue from the ErbB3 C-terminal tail (B3Y; TCTGAGCAAGGGTATGAAGAGATGAGAGCTTTTCAG) was inserted between the receptor C-terminal tail and the Halo tag.

pCMV–ErbB2–SNAP2 was generated by replacing the Halo coding sequence in pCMV–ErbB2–Halo with the SNAP coding sequence from pCMV–SNAP–D4H (Kuwashima et al., 2024; #RDB20679, RIKEN BRC), followed by introduction of the mutations required to generate SNAP2. pCMV–FKBP–ErbB1–Halo was generated by replacing the extracellular domain of ErbB1 in pCMV–EGFR(WT)–Halo with FKBP derived from FKBP–Inp54p and introducing the F36V substitution into FKBP. The plasmids generated in this study have been deposited in the RIKEN BRC Gene Bank.

### Generation of ErbB2- and ErbB4-knockout cells with CRISPR/Cas9 gene editing

Gene-editing plasmids were constructed by annealing complementary oligonucleotides (ErbB2: 5′-CACCGGACATCAATGGTGCAGATGG-3′ and 5′-AAACCCATCTGCACCATTGATGTCC-3′; ErbB4: 5′-CACCGCTGTTGCTTGAGGATAAGCA-3′ and 5′-AAACTGCTTATCCTCAAGCAACAGC-3′) and cloning them into the BbsI site of the PX459 vector (#48139, Addgene), as previously described (Ran et al., 2013). Previously generated ErbB1-knockout HeLa cells (Abe et al., 2024) or parental HeLa cells were transfected with the resulting plasmids. Following selection with 0.3 μg/mL puromycin for 3 days, individual colonies were isolated. Genomic DNA was extracted from each clone using the GenElute™ Mammalian Genomic DNA Miniprep Kit (Sigma-Aldrich). Genome editing was verified by PCR amplification using the following primer pairs: ErbB2, 5′-ACTCCTGACCCTGTCTCTGCCTTAGGTG-3′ and 5′-AGCCTCTCCTGAGTAGCTGGGACTACTG-3′; ErbB4, 5′-AGAGACCTTTTCCCCAGATACCCAAGGCA-3′ and 5′-CTCTAGCCTGGGGCAACAGAGCGAGAC-3′. PCR products were analyzed using ICE Analysis software (Synthego) to identify clones carrying the desired gene disruptions, and appropriate knockout cell lines were selected.

### Western blot analysis

Cells were lysed in ice-cold cell lysis buffer (#22352-04, Nacalai Tesque Inc.), and lysates were clarified by centrifugation at 15,000 × g for 10 min at 4°C. Protein concentrations were determined using the Pierce™ BCA Protein Assay Kit (Thermo Fisher Scientific). Equal amounts of protein were separated on 4–15% TGX precast gels (Bio-Rad Laboratories), transferred to PVDF membranes (#1704156, Bio-Rad Laboratories) using the Trans-Blot Turbo Transfer System (Bio-Rad Laboratories), and blocked with 2% non-fat dry milk in Tris-buffered saline containing 0.1% Tween-20 (TBST).

The following primary antibodies were used: anti-EGFR (1005)-G (RRID:AB_631420; #sc-03, Santa Cruz Biotechnology; 1:500), anti-phospho-EGFR (Tyr1068) (D7A5) (RRID:AB_2096270; #3777, Cell Signaling Technology; 1:1000), anti-HER2/ErbB2 (29D8) (RRID:AB_10692490; #2165, Cell Signaling Technology; 1:1000), anti-phospho-HER2/ErbB2 (Tyr1221/1222) (6B12) (RRID:AB_490899; #2243, Cell Signaling Technology; 1:1000), anti-HER3/ErbB3 (D22C5) (RRID:AB_2721919; #12708, Cell Signaling Technology; 1:1000), anti-phospho-HER3/ErbB3 (Tyr1289) (21D3) (RRID:AB_2099709; #4791, Cell Signaling Technology; 1:1000), anti-ErbB4 (C-18) (RRID:AB_2231308; #sc-283, Santa Cruz Biotechnology; 1:500), anti-phospho-HER4/ErbB4 (Tyr1284) (21A9) (RRID:AB_2099987; #4757, Cell Signaling Technology; 1:1000), anti-p44/42 MAPK (Erk1/2) (137F5) (RRID:AB_390779; #4695, Cell Signaling Technology; 1:1000), anti-phospho-p44/42 MAPK (Erk1/2) (Thr202/Tyr204) (D13.14.4E) (RRID:AB_2315112; #4370, Cell Signaling Technology; 1:1000), and anti-HaloTag polyclonal antibody (RRID:AB_713650; #9281, Promega; 1:500).

Horseradish peroxidase (HRP)-conjugated anti-rabbit IgG (RRID:AB_2099233; #7074, Cell Signaling Technology; 1:1000) was used as the secondary antibody. Immunoreactive proteins were visualized using ECL Prime Western Blotting Detection Reagent (GE Healthcare) and imaged with an ImageQuant LAS 500 imaging system (GE Healthcare).

### Single-molecule tracking (SMT) analysis

SMT analysis was performed as previously described (Kuwashima et al., 2021; Yanagawa et al., 2018; Yanagawa and Sako, 2021), with minor modifications. HeLa cells were transfected with 50 ng of plasmid DNA encoding mStayGold-, SNAP2-, or Halo-tagged proteins and cultured for at least 24 h. For imaging of SNAP2- and Halo-tagged proteins, cells were labeled with 10 nM SNAP-Cell® TMR-Star (SNAP-TMR; #S9105S, New England Biolabs) and 10 nM SaraFluor™ 650T (SF650) HaloTag ligand (#A308-01, GORYO Chemical), respectively, for 30 min in DMEM supplemented with 10% FBS. Cells were washed three times with the same medium, transferred to DMEM containing 0.2% BSA, and incubated for at least 3 h before imaging.

Single-molecule imaging was performed at 25°C in DMEM/Ham’s F-12 supplemented with HEPES and 0.2% BSA using a custom-built total internal reflection fluorescence microscope (TIRFM) based on a TiE microscope (Nikon). Fluorescent proteins were excited with a 488-nm laser (20 mW, 5% output; Sapphire 488-200, Coherent) for mStayGold, a 561-nm laser (30 mW; Sapphire 561-150, Coherent) for TMR, and a 647-nm laser (25 mW; OBIS 647, Coherent) for SF650 through a Plan Apo 100×/1.49 NA objective (Nikon) and a multiband dichroic mirror (ZT405/488/561/640, Chroma). Emission signals were separated using tandem W-View Gemini image splitters (Hamamatsu) equipped with dichroic mirrors (T550lpxr-UF2 and T650lpxr-UF2, Chroma) and emission filters (ET525/50m for mStayGold, ET590/50m for TMR, and ET700/75m for SF650; Chroma). Images (512 × 512 pixels; 65 nm/pixel) were simultaneously acquired using two ORCA Fusion BT sCMOS cameras (Hamamatsu). Time-lapse images consisting of 150 frames were recorded with an exposure time of 30 ms per frame using the Zido Auto Imaging System.

Single-molecule trajectories and variational Bayesian hidden Markov model (VB-HMM) analyses were performed using the Zido Auto Analysis System based on a two-dimensional Gaussian fitting algorithm. Subsequent analyses, including diffusion dynamics, fluorescence intensity distributions, colocalization, and statistical analyses, were performed using an updated version of smDynamicsAnalyzer, a custom Igor Pro 9.0 (WaveMetrics)-based analysis program (smDA-Igor on GitHub). Putative oligomer sizes were estimated by fitting the distribution of total fluorescence intensities with multiple Gaussian functions. Detailed analysis procedures and fitting algorithms have been described previously (Yanagawa and Sako, 2021).

Particle colocalization between A and B was calculated as described previously (Kuwashima et al., 2024). To account for differences in the expression levels of A and B, an apparent binding affinity (K_B_) was calculated. K_B_ was defined as the density of colocalized particles normalized to the product of the densities of free particles A and B.

### Depletion of PI(4,5)P_2_

Cells were transfected with Lyn_11_–FRB (#38004, Addgene) alone or together with FKBP–Inp54p (#20155, Addgene) and cultured in DMEM supplemented with 10% FBS. After incubation in DMEM containing 0.2% BSA for at least 3 h, cells were treated with 5 μM rapamycin (#sc-3504, Santa Cruz Biotechnology) for 20 min at 37°C to recruit FKBP–Inp54p to the plasma membrane and induce PI(4,5)P_2_ depletion. Cells were then transferred to 25°C and stimulated with EGF or HRG at 25°C.

### Statistical analysis

Statistical analyses were performed using Igor Pro software (WaveMetrics, Inc.) or Microsoft Excel. Experiments were repeated at least three times, as indicated in the figures and corresponding figure legends. Data are presented as the mean ± SD or mean ± SEM, as indicated in the figure legends. Statistical significance was evaluated using t-tests, as appropriate. A p value < 0.05 was considered statistically significant. Statistical significance is indicated as follows: *p* < 0.05 (*), *p* < 0.01 (**), *p* < 0.001 (***), and not significant (NS), *p* ≥ 0.05.

## Supplemental materials

**Videos 1 and 2.** Representative TIRFM images of ErbB4(WT) and ErbB4(3N), respectively. ErbB4(WT)–Halo or ErbB4(3N)–Halo was transiently expressed in ErbB2/ErbB4 DKO HeLa cells and labeled with the SF650 Halo ligand. Single ErbB4 particles were imaged by TIRFM at a temporal resolution of 30 ms for 4.5 s before HRG stimulation (left). Following stimulation with 20 nM HRG, the same cells were imaged again 5 min after HRG addition (right). Scale bar, 2 µm.

**Videos 3 and 4.** Representative TIRFM images of ErbB2(WT)/ErbB4(WT) and ErbB2(3N)/ErbB4(3N), respectively. ErbB2(WT)–SNAP2 or ErbB2(3N)–SNAP2 was coexpressed with ErbB4(WT)–Halo or ErbB4(3N)–Halo, respectively, in ErbB2/ErbB4 DKO cells at low expression levels. ErbB2–SNAP2 (green) and ErbB4–Halo (magenta) were labeled with SNAP-Cell TMR-Star and SaraFluor 650T (SF650), respectively. Single particles were imaged by TIRFM at a temporal resolution of 30 ms for 4.5 s before HRG stimulation (left). Following stimulation with 20 nM HRG, the same cells were imaged again 5 min after HRG addition (right). Scale bar, 2 µm.

## Data availability

All data generated or analyzed during this study are included in the manuscript, figures, figure supplements, and source data files.

## Acknowledgments

YS is supported by Grants-in-Aid for Scientific Research from MEXT (19H05647 and 24K01997). MA is supported by Grants-in-Aid for Scientific Research from MEXT (22K06609 and 26K09643). MY is supported by Grants-in-Aid for Scientific Research from MEXT (24K01982, 24H01266, and 25H01328) and by the Kobayashi Foundation. We thank the RIKEN BioResource Research Center for STR profiling and mycoplasma testing. We thank the Support Unit for Bio-Material Analysis, RRD, RIKEN, for DNA sequencing.

