## Supplementary figures and images for "Membrane PI(4,5)P_2_ and ErbB2 abundance regulate ErbB receptor oligomerization and kinase activation"

### Figure S1

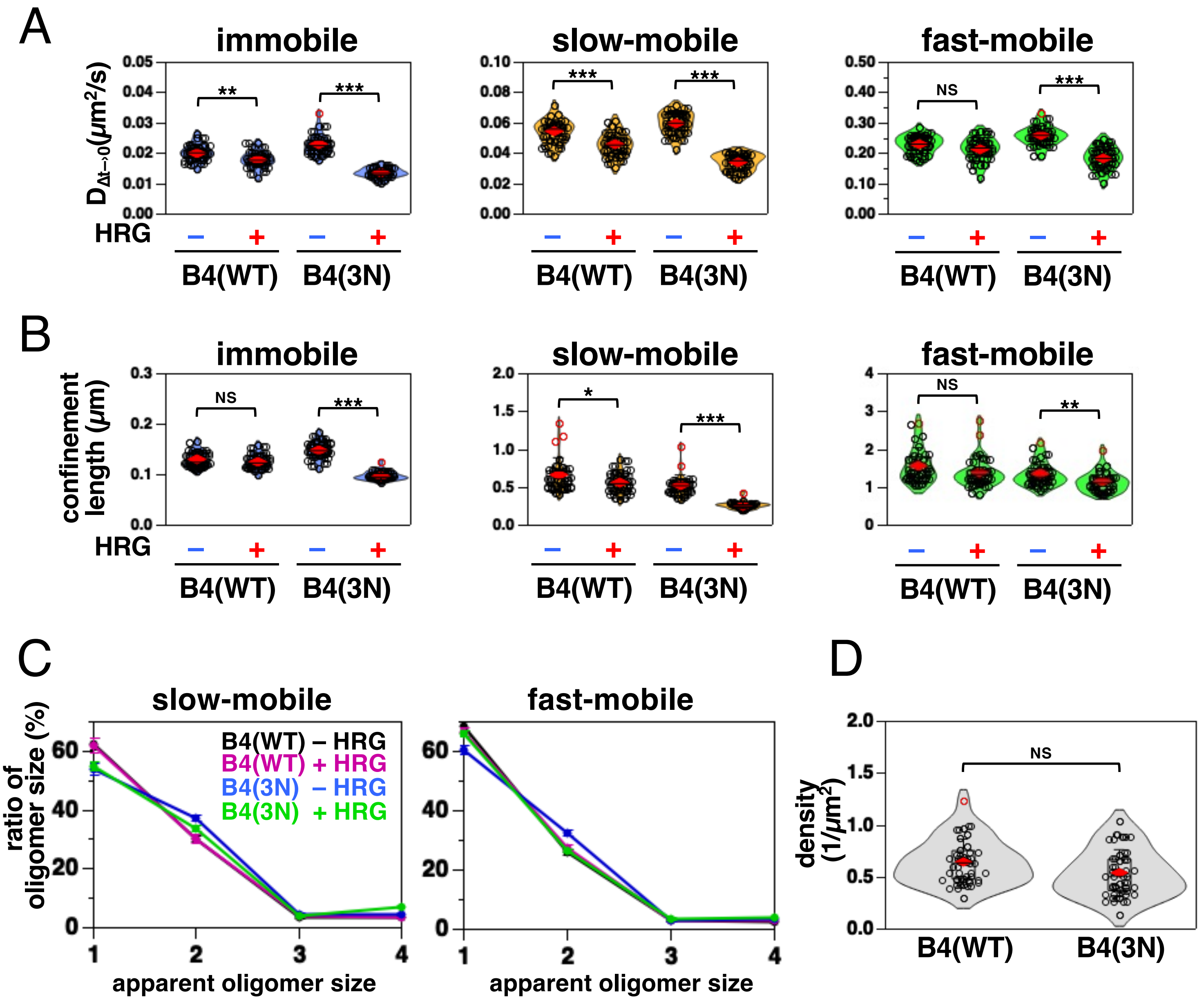

### Figure S2

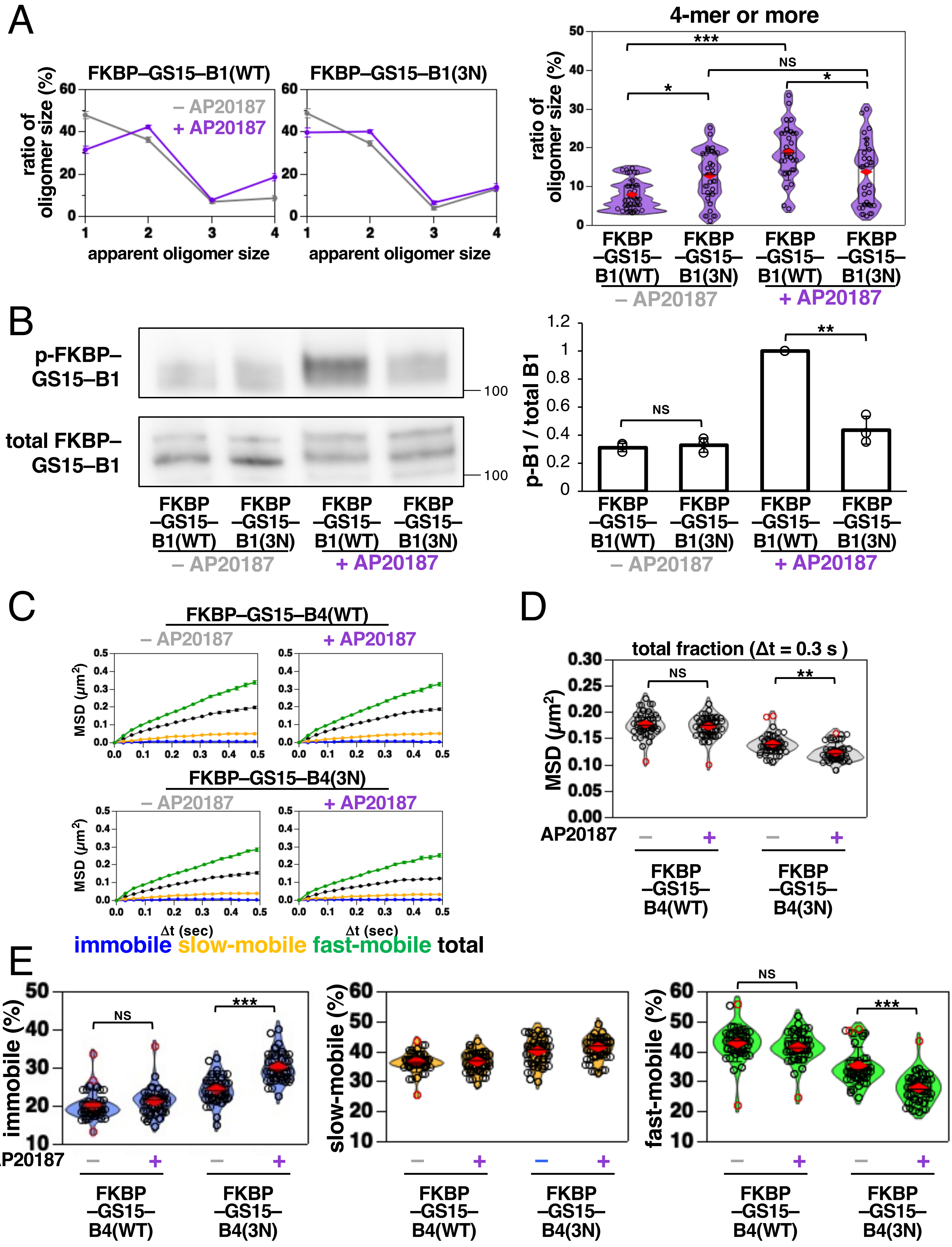

### Figure S3

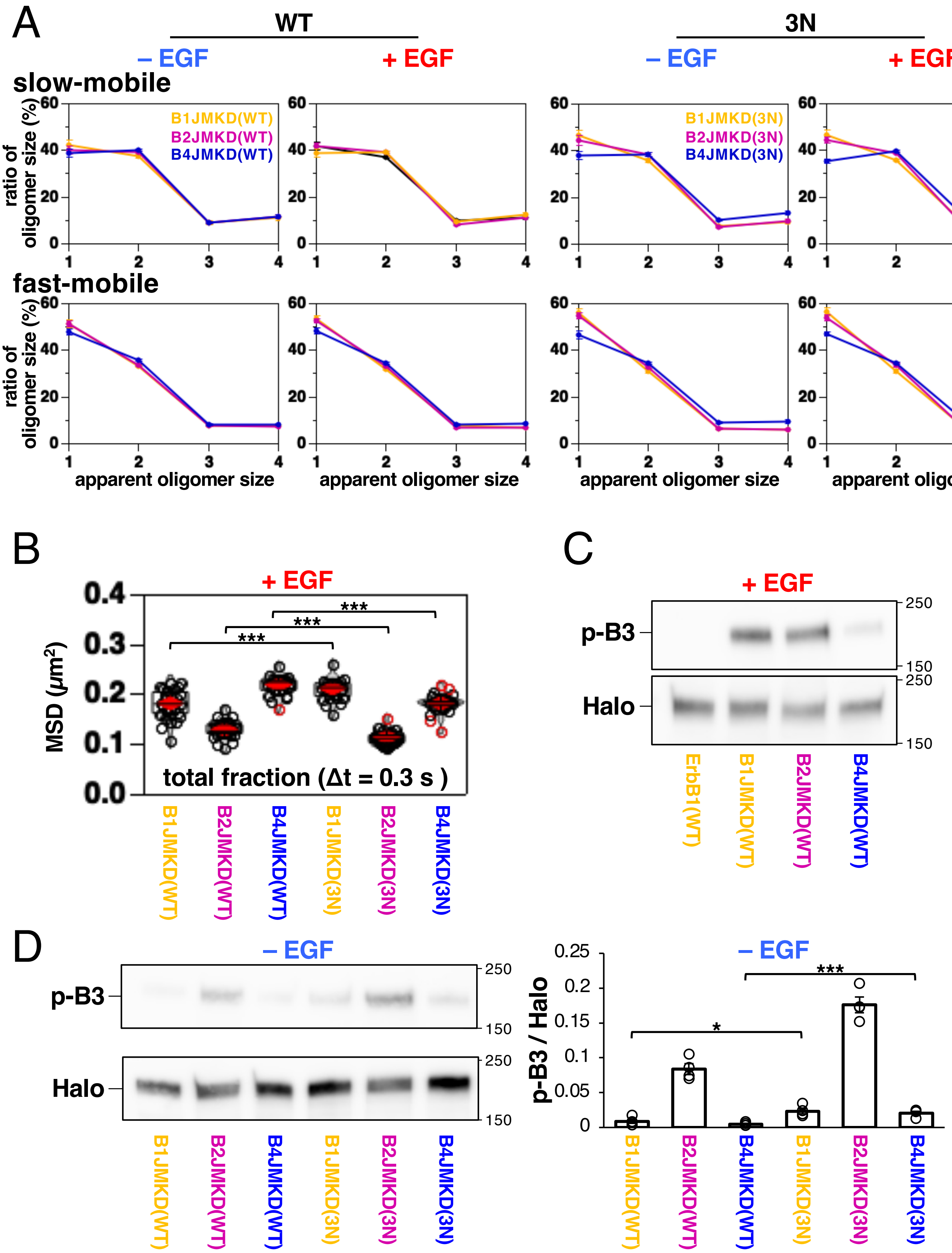

### Figure S4

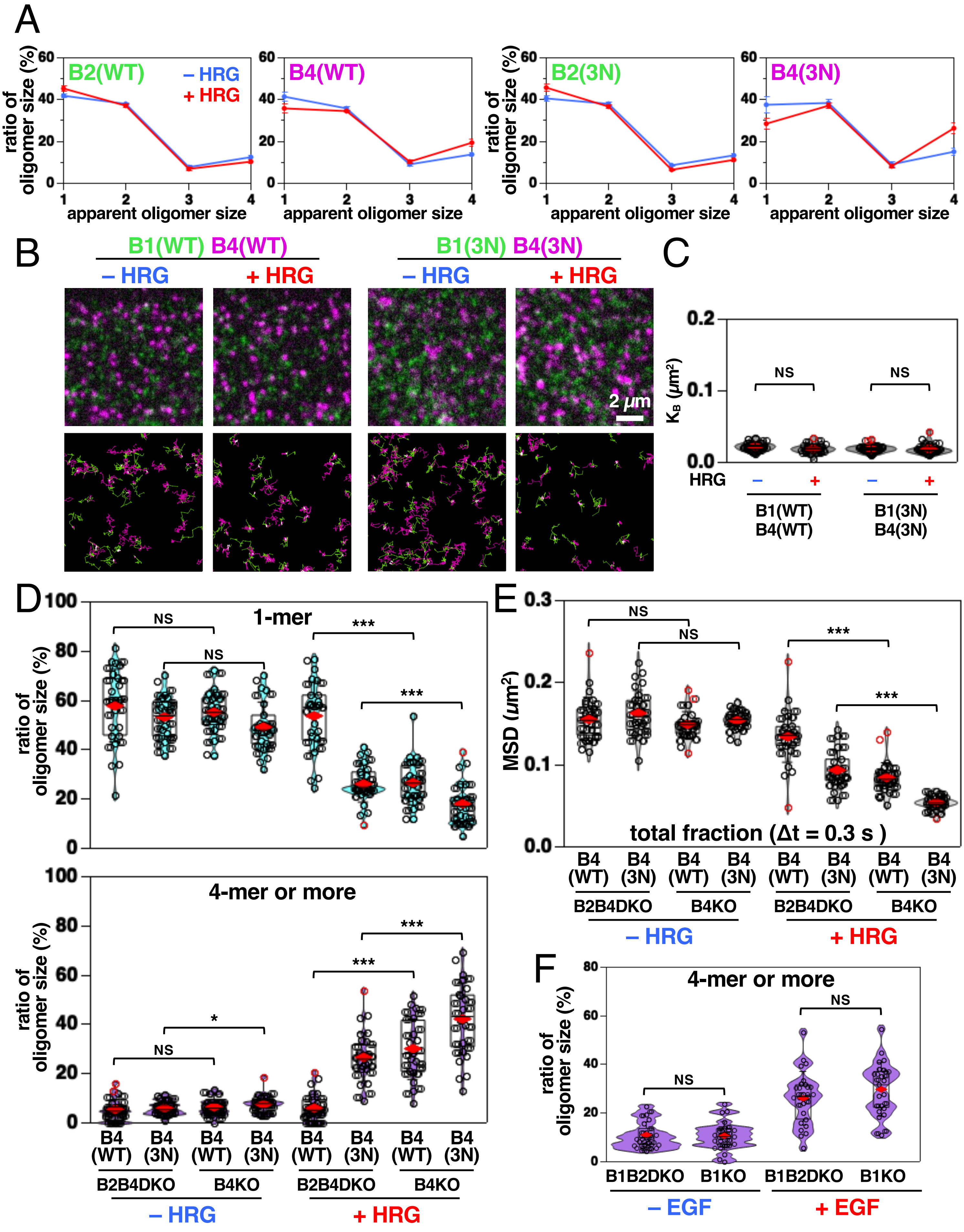

### Figure S5

Figure S5

ErbB1 JM-A

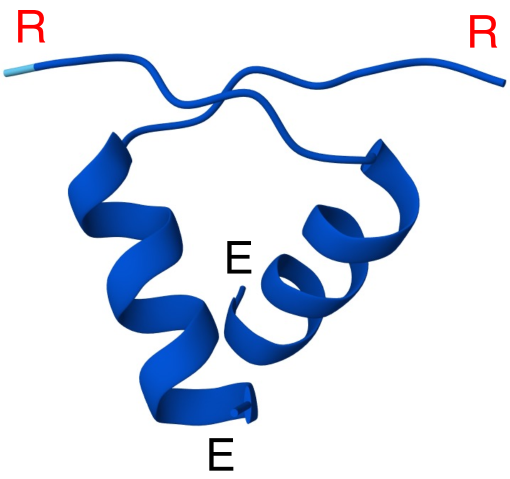


R R R H I V R K R T L R R L L Q E R E

ErbB4 JM-A

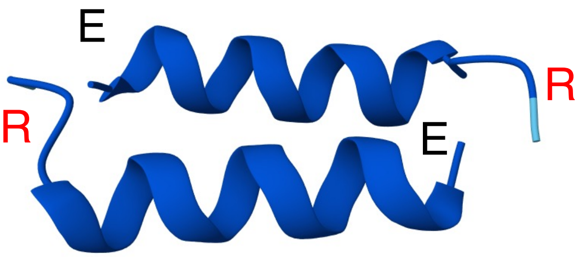


R R K S I K K K R A L R R F L E T E
